# Automated Segmentation of Kidney Nephron Structures by Deep Learning Models on Label-free Autofluorescence Microscopy for Spatial Multi-omics Data Acquisition and Mining

**DOI:** 10.1101/2021.07.16.452703

**Authors:** Nathan Heath Patterson, Ellie L. Pingry, Felipe A. Moser, Angela R.S. Kruse, Martin Dufresne, Katerina Djambazova, Allison B. Esselman, Maya Brewer, Jacqueline M. Van Ardenne, Jamie L. Allen, Amelia Wang, Mark deCaestacker, Melissa A. Farrow, Raf Van de Plas, Jeffrey M. Spraggins

**Affiliations:** Department of Biochemistry, Vanderbilt University, Nashville, TN, USA; Mass Spectrometry Research Center, Vanderbilt University, Nashville, TN, USA; Department of Cell and Developmental Biology, Vanderbilt University, Nashville, TN, USA; Delft Center for Systems and Control, Delft University of Technology, Delft, Netherlands; Department of Chemistry, Vanderbilt University, Nashville, TN, USA; Division of Nephrology and Hypertension, Department of Medicine, Vanderbilt University Medical Center, Nashville, Tennessee, USA; Department of Medicine, Vanderbilt University, Nashville, TN, USA; Department of Pathology, Microbiology and Immunology, Vanderbilt University Medical Center, Nashville, Tennessee, USA

**Author notes:** Corresponding Author* Jeffrey M. Spraggins; 465 21^st^ Ave S. Room 9160, Medical Research Building III, Vanderbilt University, Nashville, TN 37240. Aspect Analytics, C-Mine 12, 3600 Genk, Belgium (Nathan Heath Patterson). Contact emails* Nathan Heath Patterson, Ellie Pingry, Angela R.S. Kruse, Martin Dufresne, Felipe A. Moser, Katerina V. Djambazova, Allison B. Esselman, Maya Brewer, Jacqueline M. Van Ardenne, Jamie L. Allen, Mark deCaestacker, Melissa A. Farrow, Raf Van de Plas.

## Abstract

Automated spatial segmentation models can enrich spatio-molecular omics analyses by providing a link to relevant biological structures. We developed segmentation models that use label-free autofluorescence (AF) microscopy to recognize multicellular functional tissue units (FTUs) (glomerulus, proximal tubule, descending thin limb, ascending thick limb, distal tubule, and collecting duct) and gross morphological structures (cortex, outer medulla, and inner medulla) in the human kidney. Annotations were curated using highly specific multiplex immunofluorescence and transferred to co-registered AF for model training. All FTUs (except the descending thin limb) and gross kidney morphology were segmented with high accuracy: >0.85 F1-score, and Dice-Sorensen coefficients >0.80, respectively. This workflow allowed lipids, profiled by imaging mass spectrometry, to be quantitatively associated with segmented FTUs. The segmentation masks were also used to acquire spatial transcriptomics data from collecting ducts. Consistent with previous literature, we demonstrated differing transcript expression of collecting ducts in the inner and outer medulla.

## INTRODUCTION

Spatial omics analyses of tissue sections enable broad molecular interrogation of cells and multicellular structures in their native context.^1,2^ While the identification of genes, proteins, metabolites, and other molecules is inherent in spatial omics assays, their *in situ* localizations can probe the spatial relationships between molecules, cells, and multicellular structures.^3^ Compartmentalizing tissue content along biologically, anatomically, and medically relevant spatial structures is essential to discern molecular signatures specific to those structures. Thus, spatial omics data analysis can benefit from reliable, automated spatial partitioning or segmentation of data across multiple modalities. There is clear biological compartmentalization at the cell level, which aligns with the objectives of single-cell studies, but it is important to note that such an organization also occurs at larger scales. For example, at the mesoscale, many organs contain organized multicellular structures, referred to as functional tissue units (FTUs)^3^, which act in concert to fulfill biological functions.

The nephron of the mammalian kidney is divided into FTUs that include the glomerulus and 14 different tubular segments that selectively filter and excrete toxic waste products from the blood while reabsorbing water and essential solutes.^4^ In clinical pathology of the kidney, both cell and FTU morphology are examined to make diagnoses.^5–7^ For instance, glomerular sclerosis is noted by examining stained microscopy sections for cellular and extracellular perturbations of the glomerulus, such as accumulation of collagen.^8^ With the increasing presence and adoption of digital pathology^9,10^, these mesoscale FTUs have been an important automatic segmentation target for clinical analysis.^11,12^ FTUs are also gaining recognition in tissue atlas programs that seek to create spatio-molecular references, as they fit neatly along the spatial ontology between gross morphology and single cells.^13–15^ This creates an opportunity to analyze multiple cell types that act together, which in turn necessitates the development of computational workflows that consider the local environment and the molecular signal of the cells.^16–18^ For example, multiplex immunofluorescence (MxIF) and spatial transcriptomics analyses have addressed this through cell neighborhood analysis, which often recapitulates FTUs.^19^

There is clear value in linking molecular findings to distinct spatial structures in the context of tissue interrogation; however, automatic recognition and segmentation of FTUs at scale can be challenging due to a lack of appropriate training data and difficulty acquiring expert annotation.^20^ Recently, automated spatial segmentation models of the kidney have been developed using histological stains like PAS, H&E, or Masson’s Trichrome. These stains are non-specific at the FTU level and require experts for interpretation.^21^ Thus, pathologists use stain color, texture, and spatial environment to discern FTUs. As expertise is often more limited than hardware and data, annotation is a frequent bottleneck in converting scanned slide data into actionable evidence through machine learning. Furthermore, histological stains irreversibly alter the molecular content of tissues, limiting their compatibility with spatial-omics workflows. As similar workflows grow in complexity and scale, it is essential to circumvent these issues to produce robust and actionable prediction models.^22^ This can be achieved by utilizing complementary unstained microscopy to accurately recognize FTUs without compromising tissue or molecular integrity and hampering the -omics analysis. Annotation workflows utilizing ‘helper modalities’ can be developed to curate highly specific annotations by non-experts. In this manner, high-quality reliable annotations can be produced at scale and utilized by any co-registered imaging modality.

Here, we used label-free autofluorescence (AF) microscopy, which measures the fluorescence of endogenous tissue fluorophores without any stains or antibodies, to deliver automated segmentation of kidney FTUs and gross morphological structures. We demonstrated an application of this pipeline to analyze and drive downstream collection of -omics data, enabling new multimodal analyses for molecular histology. We employed MxIF microscopy as the highly specific ‘helper’ modality to computationally scale human non-expert annotations on MxIF to a much larger number of training annotations on AF. Ultimately, large training datasets were used to construct robust AF segmentation workflows, minimizing the burden on expert pathologists, data annotators, and curators. We demonstrate how these automated annotations can be used to analyze imaging mass spectrometry (IMS) measurements and to drive data collection with spatial transcriptomics.

## METHODS

### Sample curation and tissue sectioning

Normal portions of renal cancer nephrectomies from adult patients were studied. Deidentified macroscopically normal tissue samples and associated patient metadata were collected following surgery by the Vanderbilt University Medical Center Cooperative Human Tissue Network (CHTN) under the Vanderbilt University IRB protocols #181822 and #031078 (for the CHTN). Fresh frozen tissue blocks were collected from areas distant from the tumor, confirmed by stained microscopy of sequential sections. Tissues were sectioned to 10 μm thickness using a Leica cryostat (Leica Biosystems, Wetzlar, Germany) and thaw-mounted onto glass slides; indium tin oxide (ITO) coated glass slides were used for MALDI IMS experiments. A list of the samples used in this manuscript is summarized in **Table S1**.

### Autofluorescence and Multiplex Immunofluorescence data acquisition

Autofluorescence images were acquired using a Zeiss AxioScan.Z1 slide scanner (Carl Zeiss Microscopy GmbH, Oberkochen, Germany) using a 10x objective, resulting in a pixel size of 0.65 μm/px. Details on the filter cube, excitation/emission wavelengths, and exposure times can be found in **Tables S2**. Immunofluorescence microscopy was also acquired using a Zeiss AxioScan.Z1 slide scanner. A four-cycle MxIF approach was used for the immunofluorescence experiments.^2324^ Details on the immunofluorescence experiment are provided in **Table S3 and S4**. Briefly, five antibodies (Podocalyxin, Aquaporin-1, Aquaporin-2, NaCl

Cotransporter, and Uromodulin) were used to detect six FTUs (glomerulus, proximal tubule, descending thin limb, ascending thick limb, distal tubule, and collecting duct), while Collagen-IV (a1/2) was used to stain the basement membrane and indicate the border of each FTU in the kidney sample (**Figure S1**). The gross morphological regions of the kidney, namely the cortex, inner medulla, and outer medulla, were also described using these markers. Acquaporin-1 was used to discern cortex and medulla, while Uromodulin was used to distinguish the outer and inner medulla, as the TAL FTU terminates at the junction of the two medullary zones (**Figure S2**).^25^

### Whole Slide Image Registration, Annotation, and Deep-learning Segmentation Model Development

The registration and segmentation modeling workflow was conducted in four phases, as summarized in **Figure S3**. In *Phase 0*, all images for a single tissue section were collected separately and registered using *wsireg*.^26^ for cross-modality analysis. First, all MxIF images were aligned to images of the first MxIF cycle by co-registering the DAPI channels of each cycle. The max intensity projection from all non-DAPI channels was then used to register the MxIF and AF data. All images were transformed, resampled to a single coordinate space, and combined into a single pyramidal OME-TIFF^27^ file with 18 channels (AF + MxIF images). All transforms across the graph were composed and applied when writing the finalized images, resulting in a single nearest-neighbor interpolation being applied for resampling.

In *Phase 1*, kidney tissue images were manually annotated using QuPath ^28^ and its polygon annotation tool. Whole slide images were used for gross morphology annotations; both were downsampled by 4× and split into patches of 512×512p× (∼1331×1331 µm). Each patch was accompanied by a pixel-wise binary match indicating where the pixel fell within the gross morphology. For FTU annotations, small rectangular regions of approximately 30-40 tiles of size 1024×1024 pixels (665×665 μm) were manually annotated. Based on the relevant IF channels of each image, the target FTUs were manually annotated, and the resulting labels were exported using the GeoJSON ^29^ format.

In *Phase 2*, the manual annotations generated during *Phase 1* were expanded through a transfer-learning neural network approach. Using the *Detectron2* platform.^30^ These manual annotations were used to fine-tune an instance segmentation model for predicting the FTUs from the IF channels. In particular, the Mask R-CNN model with the 101 ResNet backbone was chosen for this task because of its high segmentation performance on open datasets. This was repeated separately for each FTU. The fine-tuned models were then used to generate annotations for the entire tissue section, which were, in turn, manually curated to ensure their quality, removing erroneous detections and/or improving poor detections, such as partial or missing segmentations. Through this approach, it was possible to generate a large number of high-quality annotations for each FTU, at a fraction of the time that would have been required to achieve this manually. This resulted in over 10,000 annotations for each FTU (**see Table S5**), with the exception of the glomeruli of which there are significantly fewer instances in each tissue section.

Finally, in *Phase 3*, the curated FTUs from *Phase 2* were used to train a model to predict the instance segmentations from the AF images. Similarly to the transfer-learning approach described for *Phase 2*, the *Detectron2* platform was used to fine-tune Mask R-CNN model with the 101 ResNet backbone. This was repeated separately for each FTU. For gross morphological segmentation, models that perform pixel wise semantic segmentation were used over instance segmentation models, as the gross morphological areas are not an instance of a repeating unit within the tissue. The cortex, outer medulla (inclusive of the inner and outer stripe), and inner medulla were targeted. U-nets^31^ type models were trained using the **efficientnet-b2** encoder as implemented in the **segmentation-models-pytorch** package^32^. Semantic segmentation models were developed from encoders pre-trained on *ImageNet*. ^33^ In total, 17-27 MxIF whole slide images from different tissue donors were used for training each of the 7 models of *Phase 3* (6 for FTUs and 1 for gross morphological region). These models were then tested on 7 hold-out MxIF images (**Tables S5 & S6)**.

### MALDI Imaging Mass Spectrometry

Following AF microscopy, the matrix 1,5-Diaminonapthalene was sprayed onto the tissue section using an HTX M5 Sprayer (HTX Technologies, Chapel Hill, NC). MALDI IMS data were collected in negative ionization mode using a Bruker timsTOF Flex mass spectrometer (Bruker Daltonics, Bremen, Germany) at 10 μm raster spacing.^34^ After MALDI IMS, AF microscopy was performed on the tissue section with the matrix still present to reveal laser ablation marks on the matrix layer. The image was registered to the MALDI IMS pixels using *IMS MicroLink*. ^3536^ Additional AF and MxIF microscopy was layered onto the IMS-registered image using *wsireg*. This approach provides approximately 0.25-1.50 μm accuracy (∼0.50-2.00 microscopy pixels in these images). Peak picking of the dataset was performed using *S/N* threshold of 5 on the mean spectra; peak data and total ion current (TIC) for each spectrum were extracted using the TIMSCONVERT python library^37^. Data were normalized by TIC and loaded into the R package *Cardinal*^38^. The *Cardinal* function **colocalized()** was used to compute the Pearson correlation coefficient of all IMS signals to each FTU mask. Extracted ion images and segmentations were visualized in *napari*.^39^ Putative lipid identifications (exact mass match, <5 ppm) were made with the LIPIDMAPS database.^40^ Plots were generated in *ggplot2* in R.

### Spatially-Targeted Transcriptomics Acquisition and Analysis

Tissue sections were prepared following the manufacturer’s protocol for fresh frozen tissue (GeoMx DSP Manual Slide Preparation, Nanostring Technologies, Inc., Seattle, WA). Tissues were hybridized with the Nanostring human whole transcriptome atlas probe set. Autofluorescence microscopy and region of interest (ROI) selection was performed in the Nanostring GeoMX Digital Spatial Profiler (DSP) (Nanostring Technologies, Inc., Seattle, WA). AF microscopy was acquired at 488, 550, and 647 nm, with 500 ms exposure times. Acquisition ROIs were selected based on segmentation masks from the registered AF images. Libraries were prepared for NGS following the manufacturer’s protocol (GeoMx DSP NGS readout, Nanostring Technologies, Inc., Seattle, WA). Sequencing was performed using an Illumina miniSeq with a high-output reagent cartridge.

Adapter trimming and deduplication were performed using the Nanostring NGS Data Analysis Pipeline to convert fastq files to DCC files. Analysis and visualization of the spatial transcriptomics data was performed in Nanostring GeoMX DSP software version 2.4.0.421. Probes were excluded from analysis if they failed the Grubbs outlier test in greater than 20% of ROIs or if the ratio of geometric means in all segments to within the target gene was ≤ 0.1. The limit of quantitation was calculated as 2 standard deviations above the geometric mean of the negative control probe. Q3 normalization was performed for all target groups. ROIs were excluded from analysis if their sequencing saturation was < 50%. Pathway enrichment analysis was performed requiring a minimum of 20% coverage for genes in each pathway and at least 5 genes in the pathway using 10,000 permutations.

## RESULTS

### Performance of AF-based models for gross morphological and FTU segmentation

Performance for gross morphological segmentation, shown for one example in **Figure 1a**, was computed using the Dice-Sorensen Coefficient (DSC), which measures the degree of overlap between the curated ground truth annotations and the models’ predictions in a pixel-wise fashion. DSC values of 0 indicate the model did not detect the structure (false negative), and 1 indicates a perfect match between the model’s segmentation pixels and the ground truth annotation pixels. Here, gross morphological segmentation probability maps were binarized using a threshold of 0.40. **Figure 1b** shows an overlay of gross morphological segmentation results (filled color) and ground truth annotations (polygons). The mean DSC for cortex, inner medulla, and outer medulla across the 8 test whole-slide images was >0.85 (**Figure 1c**).

**Figure 1.**
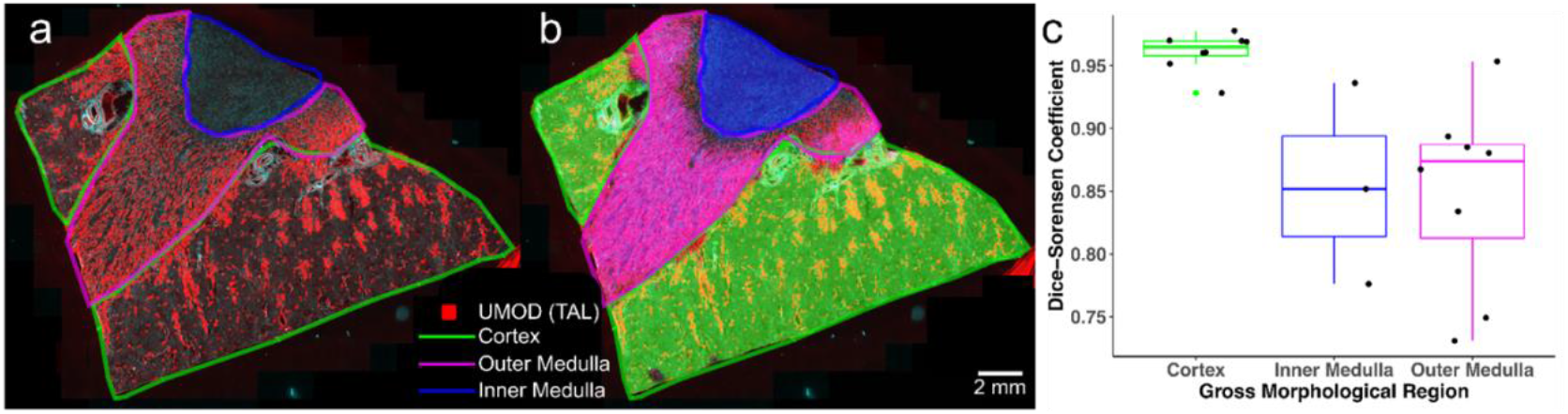
Gross morphological segmentation analysis and results. **(a)** Visualization of Uromodulin, a marker for thick ascending limb (red) alongside ground truth annotations of kidney cortex (green), kidney outer medulla (magenta), and kidney inner medulla (blue). Uromodulin is displayed because thick ascending limb starts in the outer medulla and is not present in the inner medulla. **(b)** Overlay of gross morphological segmentation results (filled color) and ground truth annotations (polygons). **(c)** Plot of Dice-Sorensen Coefficient of whole slide autofluorescence images gross morphological segmentation. UMOD = Uromodulin. TAL = Thick ascending limb.

FTU segmentation performance was evaluated at the whole-slide image level. An overlay of the FTU segmentation masks and AF microscopy image can be seen in **Figure 2a. Figure 2b** shows an overlay image of the predicted and ground-truth proximal tubules. A global DSC was used to determine the absolute pixel-wise segmentation accuracy across the test set of 8 whole-slide images, showing a mean DSC of ∼0.90 for all FTUs except the descending thin limb (**Figure 2c**). Additionally, instance-level metrics, including F1-score, precision (the positive predictive value), and recall (true positive rate), were evaluated for each structure (**Figure 2e-g**). Each metric reports a value between 0 and 1, indicating the agreement between the trained model and the ground truth annotation. Precision and recall metrics capture the number of correct detections (true positives), incorrect detections (false positives), and missing detections (false negatives). Here, segmentations with a DSC ≥ 0.75 were treated as true positives. In total, the instance-level F1-score closely resembled the DSC coefficient, precision was > 0.90 for all structures, and recall was ∼0.80-0.85 for each structure except the descending thin limb. Supplemental **Table S6** reports the count statistics for the test set FTUs, and **Tables S7** and **S8** report metric values and summary statistics across all samples.

**Figure 2.**
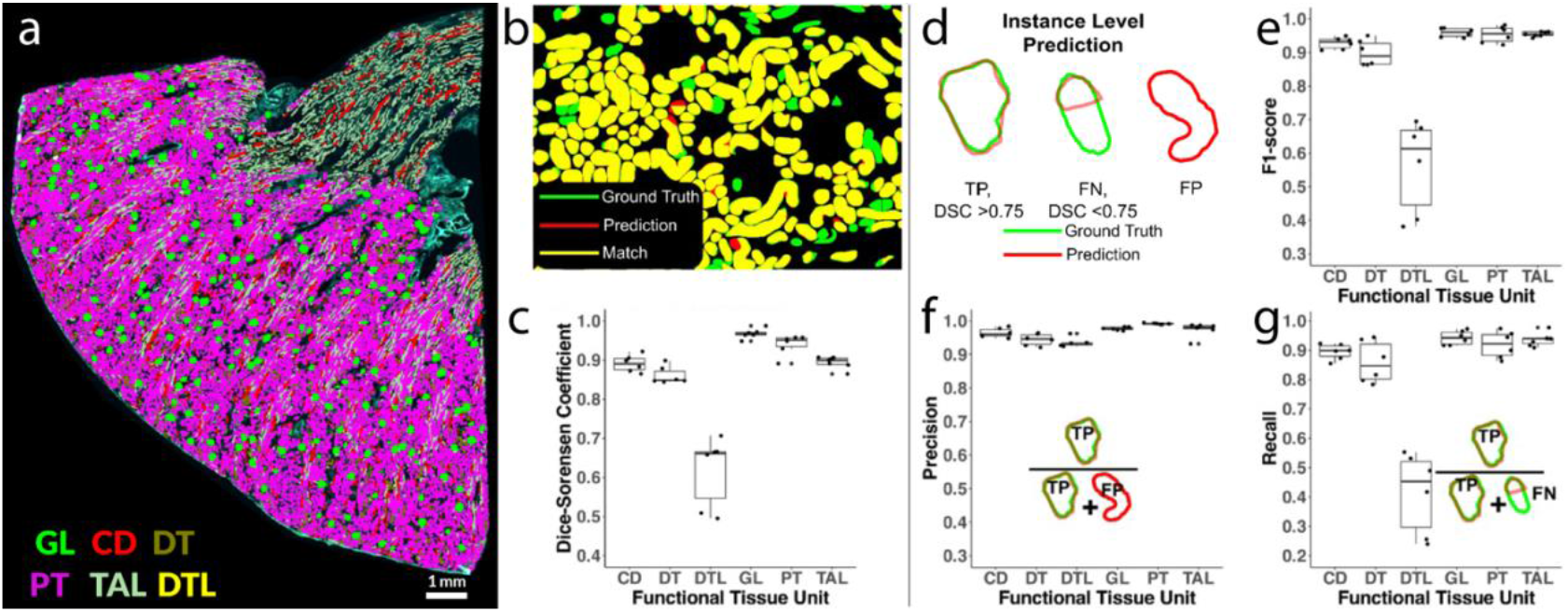
Global pixel-wise and instance-level accuracy analysis of whole slide segmentation results for autofluorescence (AF) based functional tissue unit segmentation (FTU) of an independent test set. **(a)** Whole slide view of an AF image with overlaid filled polygons for each segmented FTU. **(b)** Overlay of binary mask of proximal tubule segmentation and ground truth proximal tubule annotation from kidney cortex. **(c)** Global metric Dice-Sorensen coefficient calculated from the test images. **(d)** Visual demonstration of local instance level true positives (TP), false negatives (FN), and false positives(FPs) are called for each instance of an FTU. **(e)** Plot of F1-score for the instance level metrics for each functional tissue FTU. **(f)** Plot of the precision of the model at the instance level for each FTU and diagram of how precision is calculated. **(g)** Plot of the recall of the model for each FTU at the instance level and diagram of how recall is calculated. *GL – Glomerulus, CD - Collecting Duct, DT - Distal Tubule, PT - Proximal Tubule, TAL - Thick Ascending Limb, DTL - Descending Thin Limb*

### Analysis of molecular imaging by MALDI IMS using automated label-free FTU Segmentation

Automated AF-based segmentation maps can enrich MALDI IMS data analysis by providing dense pixel labels of each FTU. **Figure 3a** shows the whole-slide image prediction of FTUs and the region of interest for IMS data collection, which included both cortex (left) and medulla (right). The AF and IMS data were registered to intersect the instance segmentations with the IMS data, providing a list of IMS pixels for each FTU. Their degree of intersection (0-1) was calculated using the physical coordinate space, which entails mapping each IMS pixel’s bounding box and each FTU’s polygon coordinates to μm. Compared to rasterized approaches, where each image is sampled to the same pixel dimension, this approach can mitigate warping effects due to the mismatch in pixel spacing between microscopy (0.65 μm/px) and IMS (10 μm/px) data. The Pearson Correlation Coefficients between all IMS peaks and FTU segmentation masks were calculated; the top MALDI IMS lipid signals associated with each FTU are shown in **Figure 3b. Figure 3c** shows the spatial distributions of the top-ranking IMS signals for four FTUs and overlays of the FTU mask and IMS signal. The lower correlation values indicate that many species are not specific to a single FTU. Indeed, most IMS signals are present in multiple structures (**Figure 3c**). For example, IMS signals associated with the collecting duct, distal tubule, and glomerulus not only show a distinct signal in those FTUs, but also localize to other structures in the sampled area.

**Figure 3.**
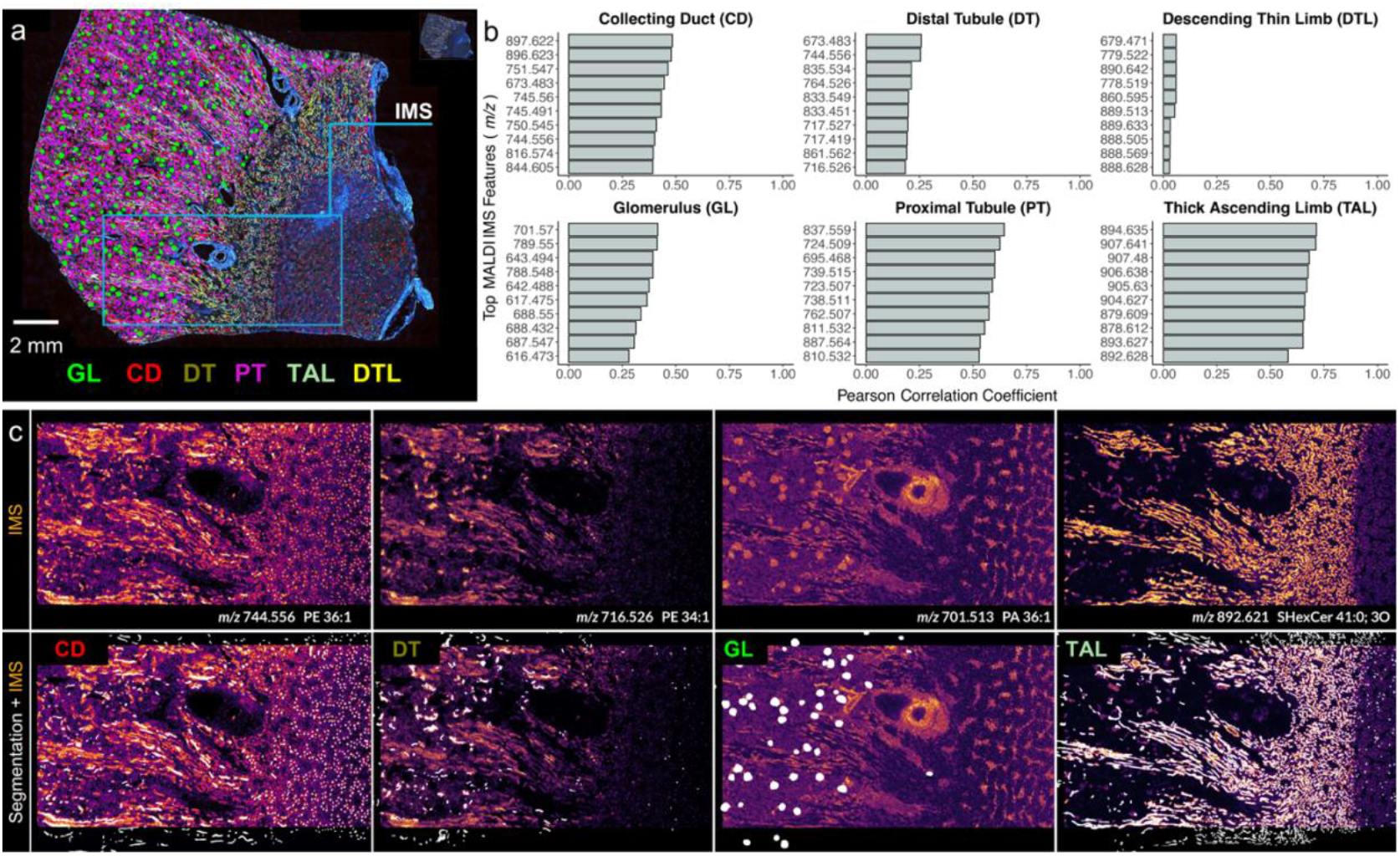
Analysis of matrix-assisted laser desorption/ionization imaging mass spectrometry (MALDI IMS) data by functional tissue unit (FTU) autofluorescence (AF) segmentation masks generated on the same tissue section. **(a)** Whole slide AF image with FTU segmentations of different colors representing different functional tissue units. The MALDI IMS region of interest for data acquisition is highlighted in cyan. **(b)** Pearson’s correlation coefficient plots of FTU masks and IMS signals. Calculations were performed after registration of MALDI IMS with AF microscopy and intersection of microscopy FTU masks with IMS pixels. **(c)** Top ranking MALDI IMS signals for 4 FTUs and overlay of FTU mask (white) with MALDI IMS signal. *GL – Glomerulus, CD - Collecting Duct, DT - Distal Tubule, PT - Proximal Tubule, TAL - Thick Ascending Limb, DTL - Descending Thin Limb*

### Spatially targeted transcriptomic analysis guided by automated label-free segmentation

Without compromising tissue or molecular integrity, label-free AF can provide segmentation maps, which can be used for spatially targeted data collection. Here, both gross morphological and FTU level instance segmentations (**Figure 4a**) were integrated to select collecting ducts from the inner and outer medulla, as shown in **Figure 4b and 4c**, respectively. These segmentation maps were used for data collection by the GeoMX Digital Spatial Profiler platform; the computational workflow can be seen in supplemental **Figure S4**. After sequencing, the ROIs were quality controlled: 6 regions passed, including 3 inner and 3 outer medulla collecting duct areas. **Figure 4d** shows a volcano plot of transcript targets and specifically highlights AQP6 as being more expressed in the outer medulla. Consistent with previous literature, we found that AQP6 was not abundantly expressed in the collecting ducts of the inner medulla.^41^ These results provide simultaneous validation of the gross morphological and the FTU level segmentations. **Figure S5** highlights boxplots of all AQP targets detected, showing AQP2 and AQP3 expression was universally high across the collecting ducts of the inner and outer medulla, as expected. **Figure 4e** shows the pathway analysis of collecting ducts from the inner and outer medulla, revealing statistically significant (p = 0.0087) differential respiratory electron transport and TCA cycle pathways. This analysis yielded nearly 70% gene coverage; **Figure 4f** shows the differential Z-scores for the top 64 differential markers with three major gene families represented: ATP, COX, and NDUF.

**Figure 4.**
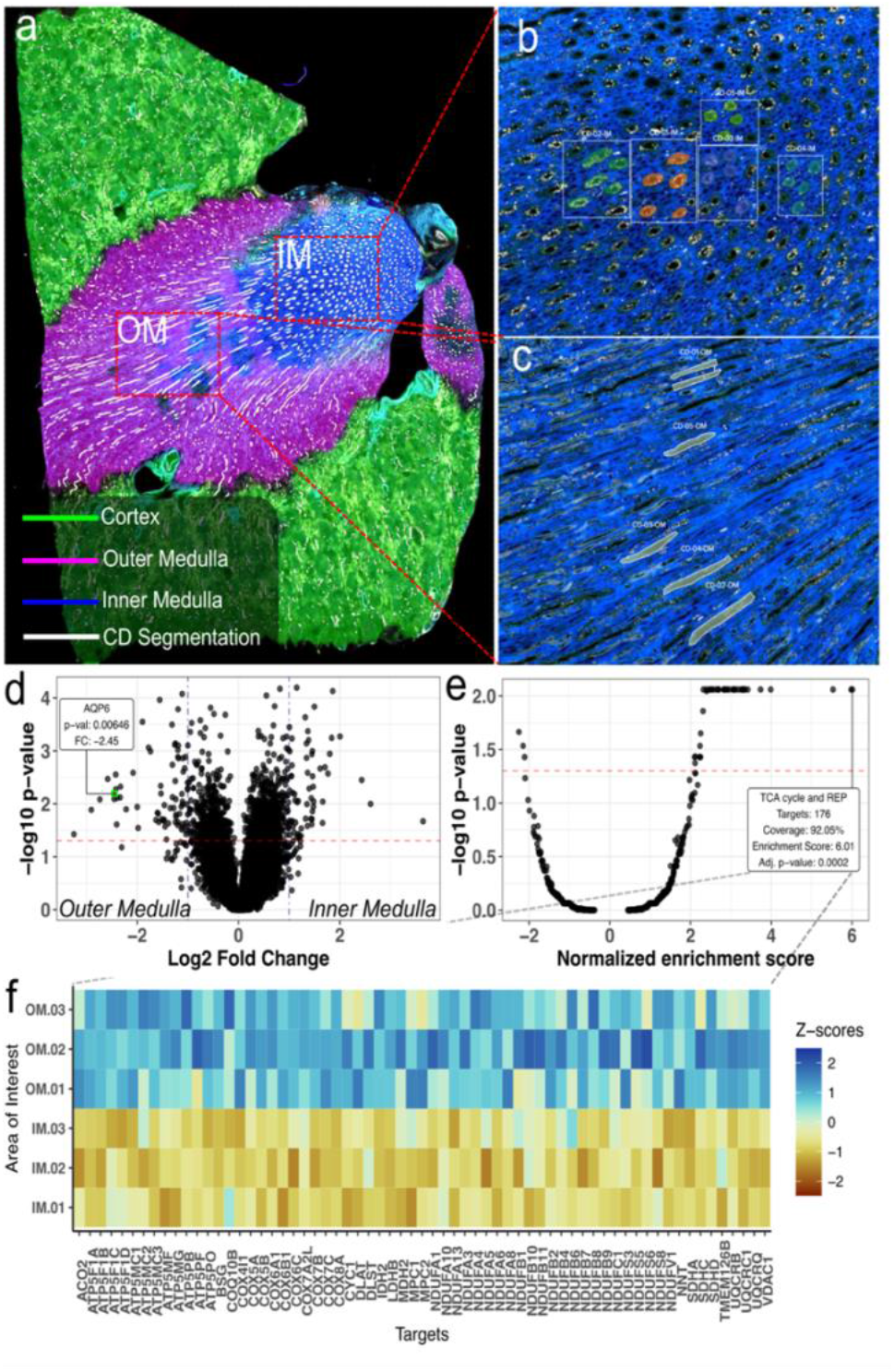
Spatially targeted transcriptomics of the collecting duct (CD) in the inner and outer medulla of a human kidney tissue section after AF-based functional tissue unit (FTU) and gross morphological whole slide image segmentation. (a) Overview of the whole slide image with gross morphological areas and CDs segmented. (b) High magnification view of the inner medulla region and the five areas of spatial transcriptomic data collection for the CD within inner medulla highlighted in different colors. (c) High magnification view of the outer medulla and the five CD ROIs in this region highlighted in green. (d) Volcano plot of Log2 fold change and -log10 p-values between transcripts detected in the inner medullary CDs (left) and outer medullary CDs (right) with the positive class being the inner medulla. Highlighted relative expression and confidence scoring for AQP6 is shown in panel inset. (e) Pathway enrichment scores for the comparison between collecting ducts in inner and outer medulla. Inset highlights the TCA cycle and respiratory electron transport pathway (REP) being enriched. (f) Heatmap of the top 64 differential transcripts in the inner and outer medulla from the TCA cycle and respiratory electron transport pathway.

## DISCUSSION

### Use of label-free autofluorescence for direct image analysis tasks and limitations to the modeling approach

AF microscopy provides non-specific images of endogenous fluorophores without sample preparation, making it highly suitable for integration with spatial omics assays. Previous studies have performed virtual staining by using AF images to predict histological stains, which have convinced blinded pathologists.^42^ Pseudo-staining approaches can provide crucial diagnostic information without the labor and cost associated with histological staining procedures. However, they still require interpretation by trained pathologists, which ultimately places a bottleneck on automated analyses. Alternatively, generating segmentations as unambiguous outputs can be more practical for spatial omics workflows. In a recent example, segmentation workflows of virtual stains were used to predict IF and cell segmentation masks.^43,44^ These studies suggested that with adequate data curation and modeling, image analysis tasks (classification, segmentation, etc.) could be performed directly from label-free data, foregoing the need for transformation to virtual stains. Building on these studies, we used AF microscopy to accurately segment multiple FTUs in human kidney whole-slide images and to generate masks for downstream spatial omics data collection and analyses. Training data were developed using MxIF to label FTUs, which were transferred to the AF microscopy. Ultimately, the models segment ∼80% of the kidney tissue into 6 FTUs based on antibody labels (**Figure 2a**). In contrast to other whole-slide image segmentation workflows, these analyses were conducted on fresh-frozen samples that do not undergo chemical fixation, which affects the molecular analysis. However, we posit adapting these models to use autofluorescence from FFPE (popular in clinical settings) would not be challenging, as it has already been demonstrated for virtual staining tasks.^45,46^

### Competitive performance in segmentation across gross morphological and FTU spatial scales

We developed a large-area segmentation model to discern gross kidney tissue morphology, including the cortex, outer medulla, and inner medulla. These regions provide valuable high-level information about the proportions of each gross morphology and are valuable for spatial atlases that seek to go from whole organ to single cell. Segmentation was performed using a U-net model that produces pixel-wise labels of the whole-slide image analogous to the modeling found in the landmark digital pathology CAMELYON challenge.^47^ While appropriate for discerning gross morphological regions, this approach is not ideal for detecting repetitive multicellular structural units like the FTUs of the nephron. Therefore, an instance segmentation model, Mask R-CNN, was used to individually segment each FTU in the section, permitting analysis of each discrete structure. We created separate models for each FTU, decreasing the time needed to curate the data necessary for training a performant model. Creating separate models was also advantageous due to the relative density of different FTUs in the tissue. As shown in supplemental **Tables S5** and **S6**, there were far more instances of proximal tubules than glomeruli, for example. Therefore, to train a model that is inclusive of all FTUs, a much greater number of proximal tubules would need to be curated. All training areas would need to be labeled to avoid the model training on images containing unlabeled areas with known FTUs. This remains a limitation of the current study, as a comparison of all-FTU models vs per-FTU models was not made.

Segmentation performance for gross morphology used DSC to evaluate the overlap between human annotation and machine segmentation. The cortex performed well (mean DSC >0.95), but the inner and outer medulla were more challenging (mean DSC <0.90) to segment. Defining the transition zone from the inner to the outer medulla precisely is difficult even in the ground truth data, as there are no explicit markers for the two regions. Furthermore, most training samples contained cortex and medulla but only 10 of the 26 contained both inner and outer medulla, limiting the available training data and affecting performance. Ultimately, this transition region was segmented as neither, as seen in **Figure 1**. Despite lower performance, the models provided actionable information as shown in the spatial transcriptomics workflow.

FTU segmentation evaluation was performed in two ways. First, global DSC indicates pixel-wise performance across the entire evaluated region for each FTU. Second, instance-level metrics were computed by first filtering structures that were well-segmented (DSC > 0.75) and treating them as true positives (**Figure 2d**). In this metric, inaccurate instance segmentation was less harshly penalized, and the segmentation was actionable for spatial omics analysis. Our results show high precision (mean precision > 0.95 for all structures) with lower recall (mean recall 0.90 for all structures except descending thin limb at 0.50). These results indicate that the model’s predictions are very accurate (high precision, few false positives), but not always comprehensive (lower recall, more false negatives). While additional performance gains would be valuable, the obtained few false positives, high recall, and high global DSC values were sufficient for reliable downstream data analysis.

Our workflows performed similarly to other studies using histological stains for functional tissue unit segmentation.^11^ For example, Jayapandin et al. reported delineation of proximal and distal tubules with 0.83 and 0.89 accuracy, respectively.^12^ Because our model uses MxIF with spatially specific markers, it can provide higher structural specificity. For example, we can further subdivide the tissue into proximal tubules, descending thin limb, thick ascending limb, distal tubules, and collecting ducts. Each of these subdivisions relays additional information and expands the scope of possible downstream analyses. On the other hand, our model did not perform as strongly for the segmentation of glomeruli. This is likely due to the glomerulus being weakly autofluorescent and thus less distinct against the background (**Fig 1g**). Additionally, our study did not subdivide FTUs with evidence of disease. This could be achieved in future studies by incorporating antibodies that mark features impacted by disease (*i*.*e*., sclerotic glomeruli) to re-train and optimize segmentation models.

### Automated molecular profiling of FTUs from MALDI IMS

AF is advantageous to guide MALDI analysis because it is performed immediately following tissue sectioning. Therefore, the tissue is not subjected to chemical washes, MALDI matrix deposition, or on-tissue laser ablation, which can change the appearance of histological stains and affect tissue integrity. Our approach combines the AF and MALDI IMS to analyze molecular data (*i*.*e*., ion images) by FTUs (**Figure 3**). One challenge of this approach is that MALDI IMS and microscopy have a significant spatial resolution gap. IMS is routinely performed at 10 μm spatial resolution, whereas microscopy has an estimated 15x higher resolution (0.65 µm). Therefore, it is challenging to analyze very small structures, such as FTUs, that may be represented by only 10s of IMS pixels. To work with the multiscale nature of the data, we used data registration^35^ and IMS pixel-structure deconvolution approaches^48,49^, which excluded IMS pixels that overlapped less than 80% with the FTU.

Ultimately, this analysis provided visual co-localization of lipid ions to specific FTUs. The degree of intersection of an IMS signal and a particular FTU was computed using Pearson correlation coefficients. Overall, these values were lower than expected (top signals ≤ 0.75), as one input is a binary vector indicating FTU (or not FTU) and the other is a continuously varying IMS signal. Some ions showed exclusive and unmistakable colocalization with the FTU of interest, as seen for ion *m/z* 892.621 (SHexCer 41:0; 3O) in the thick ascending limb (**Figure 3c)**. More commonly, however, ions were not unique to a specific FTU. For example, in **Figure 3c**, PE (34:1) (*m/z* 716.526) was present in the distal tubules, but was also found in other areas, reducing the correlation coefficient. This challenge can be compounded by the dynamic range of MALDI IMS experiments, where a small band of intensity may belong to distal tubules, but higher intensities are partitioned to other FTUs. Furthermore, lower correlation coefficients are unsurprising given that MALDI IMS is label-free and readily detects lipids that form important membrane structures, which exist in multiple FTUs. This proof-of-concept work can be further enhanced by integrating advanced approaches for cohort analysis ^50^ or those intended to properly subdivide IMS signal bands into distinct spatial regions^51^.

### Recapitulating known biology of kidney collecting ducts and differential transcriptomic analysis of inner and outer medulla collecting ducts enabled by segmentation and automated spatial targeting

With many spatial transcriptomics experiments using microscopy to guide acquisition or analysis, methods that apply to label-free microscopy are well positioned to impact the burgeoning field. In our analysis, we use the hierarchical segmentation data to nest collecting duct segmentations in the outer and inner medulla. The segmentations were used as regions for targeted data acquisition. Differential analysis through t-tests of 3 replicates on inner and outer medulla showed known biology and interesting effects around cellular energy processing. We found AQP2 and AQP3 highly expressed in both inner and outer medulla. We observed AQP6 to have a 2.45 Log2 fold change in the outer medulla versus the inner medulla, recapitulating known kidney aquaporin biology. Pathway analysis revealed potential upregulation of mitochondrial energy pathways related to the tricarboxylic acid (TCA) cycle with statistical significance and high coverage. The medullary collecting duct, specifically the outer medulla segment, is populated with mitochondria-rich intercalated cells. Enrichment of TCA cycle transcripts in the outer relative to the inner medulla is consistent with the abundance of intercalated cells in the outer medulla^52^. Interestingly, intercalated cells express high levels of OXGR1, the receptor for 2-oxoglutarate, a TCA cycle intermediate. OXGR1 and 2-oxoglutarate are part of a paracrine signaling loop between the proximal and distal nephron that regulates salt reabsorption when acid-base homeostasis is perturbed^53^. The precision of the collecting duct segmentation approach allows for acquisition of multi-omic measurements of medullary intercalated cells, thus potentially providing insight into the mechanism of intrarenal paracrine signaling events. Understanding the cross-talk between proximal tubules and intercalated cells of the outer medullary collecting duct during disease states is key to addressing acid-base homeostasis as well as the regulation of fluid and electrolyte balance during stressful conditions. Leveraging the high spatial specificity of this workflow permits investigation of particular cell types or FTU across the tissue with robust spatial resolution, providing a foundation for comprehensive analysis of renal regulation.

## Acknowledgements

This work was supported by: the National Institutes of Health (NIH) Common Fund and National Institute of Diabetes And Digestive And Kidney Diseases (NIDDK) under Award Numbers U54DK134302, U54DK120058, and U01DK133766 (J.M.S. and R.V.); National Institute Of Allergy And Infectious Diseases (NIAID) under Award Numbers R01AI138581 (J.M.S. and R.V.); the NIH National Institute On Aging (NIA) under Award Number R01AG078803 (J.M.S. and R.V.); National Eye Institute U54EY032442 (J.M.S. and R.V.); the National Science Foundation Major Research Instrument Program CBET – 1828299 (J.M.S.); Chan-Zuckerberg Initiative grants 2021-240339 and 2022-309518 (L.G.M. and R.V.). K.V.D was supported by NIDDK training grant (T32DK007569-34).

## References

1. Moffitt, J. R., Lundberg, E. & Heyn, H. The emerging landscape of spatial profiling technologies. Nat Rev Genet (2022) doi:10.1038/S41576-022-00515-3.

2. Lewis, S. M. et al. Spatial omics and multiplexed imaging to explore cancer biology. Nature Methods 2021 18:9 18, 997–1012 (2021).

3. de Bono, B., Grenon, P., Baldock, R. & Hunter, P. Functional tissue units and their primary tissue motifs in multi-scale physiology. J Biomed Semantics 4, 22 (2013).

4. Knepper, M. & Burg, M. Organization of nephron function. Am J Physiol 244, (1983).

5. Barisoni, L. et al. Digital pathology evaluation in the multicenter nephrotic syndrome study network (NEPTUNE). Clinical Journal of the American Society of Nephrology 8, 1449–1459 (2013).

6. Mariani, L. H., Martini, S., Barisoni, L. & et al. Interstitial fibrosis scored on whole-slide digital imaging of kidney biopsies is a predictor of outcome in proteinuric glomerulopathies. Nephrol Dial Transplant 33, 310–318 (2018).

7. Schelling, J. R. Tubular atrophy in the pathogenesis of chronic kidney disease progression. Pediatr Nephrol Berl Ger 31, 693–706 (2016).

8. Muda, A. O., Ruzzi, L., Bernardini, S., Teti, A. & Faraggiana, T. Collagen VII expression in glomerular sclerosis. J Pathol 195, 383–390 (2001).

9. Baxi, V., Edwards, R., Montalto, M. & Saha, S. Digital pathology and artificial intelligence in translational medicine and clinical practice. Modern Pathology 2021 35:1 35, 23–32 (2021).

10. Barisoni, L., Gimpel, C., Kain, R. & et al. Digital pathology imaging as a novel platform for standardization and globalization of quantitative nephropathology. Clin Kidney J 10, 176–187 (2017).

11. Bouteldja, N. et al. Deep Learning-Based Segmentation and Quantification in Experimental Kidney Histopathology. J Am Soc Nephrol 32, 52–68 (2020).

12. Jayapandian, C. P. et al. Development and evaluation of deep learning–based segmentation of histologic structures in the kidney cortex with multiple histologic stains. Kidney Int 99, 86–101 (2021).

13. Consortium, H. The human body at cellular resolution: the NIH Human Biomolecular Atlas Program. Nature 2019 574:7777 574, 187–192 (2019).

14. de Boer, I. H. et al. Rationale and design of the Kidney Precision Medicine Project. Kidney Int 99, 498–510 (2021).

15. Börner, K. et al. Anatomical structures, cell types and biomarkers of the Human Reference Atlas. Nature Cell Biology 2021 23:11 23, 1117–1128 (2021).

16. Hansen, J. et al. A reference tissue atlas for the human kidney. Sci Adv 8, 4965 (2022).

17. Parra, E. R. Methods to Determine and Analyze the Cellular Spatial Distribution Extracted From Multiplex Immunofluorescence Data to Understand the Tumor Microenvironment. Front Mol Biosci 8, 575 (2021).

18. Satija, R., Farrell, J. A., Gennert, D., Schier, A. F. & Regev, A. Spatial reconstruction of single-cell gene expression data. Nature Biotechnology 2015 33:5 33, 495–502 (2015).

19. Palla, G. et al. Squidpy: a scalable framework for spatial omics analysis. Nature Methods 2022 19:2 19, 171–178 (2022).

20. Reza Tizhoosh, H. & Pantanowitz, L. Artificial Intelligence and Digital Pathology: Challenges and Opportunities. J Pathol Inform 9, (2018).

21. Wahab, N. et al. Semantic annotation for computational pathology: multidisciplinary experience and best practice recommendations. J Pathol Clin Res 8, 116–128 (2022).

22. Marini, N. et al. Unleashing the potential of digital pathology data by training computer-aided diagnosis models without human annotations. npj Digital Medicine 2022 5:1 5, 1–18 (2022).

23. Brewer, M. et al. Multiplex Immunofluorescence on Fresh Frozen Tissue-V2. (2022) doi:10.17504/PROTOCOLS.IO.BS68NHHW.

24. Neumann, E. K. et al. Protocol for multimodal analysis of human kidney tissue by imaging mass spectrometry and CODEX multiplexed immunofluorescence. STAR Protoc 2, 100747 (2021).

25. Chen, L. et al. Renal-tubule epithelial cell nomenclature for single-cell rna-sequencing studies. Journal of the American Society of Nephrology 30, 1358–1364 (2019).

26. Patterson, H. & Manz, T. NHPatterson/wsireg: wsireg v0.3.5. (2022) doi:10.5281/ZENODO.6561996.

27. Besson, S. et al. Bringing Open Data to Whole Slide Imaging. Lecture Notes in Computer Science (including subseries Lecture Notes in Artificial Intelligence and Lecture Notes in Bioinformatics) 11435 LNCS, 3–10 (2019).

28. Bankhead, P. et al. QuPath: Open source software for digital pathology image analysis. Scientific Reports 2017 7:1 7, 1–7 (2017).

29. Butler, H. et al. The GeoJSON Format. (2016) doi:10.17487/RFC7946.

30. Wu, Y., Kirillov, A., Massa, F., Lo, W.-Y. Lo & Girshick, R. Detectron2. https://github.com/facebookresearch/detectron2 (2019).

31. Ronneberger, O., Fischer, P. & Brox, T. U-Net: Convolutional networks for biomedical image segmentation. Medical image Computing and Computer-Assisted Intervention – MICCAI 2015 9351, 234–241.

32. Iakubovskii, P. Segmentation Models Pytorch. GitHub repository (2019).

33. Deng, J. et al. ImageNet: A large-scale hierarchical image database. 248–255 (2010) doi:10.1109/CVPR.2009.5206848.

34. Spraggins, J. M. et al. High-Performance Molecular Imaging with MALDI Trapped Ion-Mobility Time-of-Flight (timsTOF) Mass Spectrometry. Anal. Chem. 91, 14552–14560 (2019).

35. Patterson, N. H., Tuck, M., van de Plas, R. & Caprioli, R. M. Advanced Registration and Analysis of MALDI Imaging Mass Spectrometry Measurements through Autofluorescence Microscopy. Anal Chem 90, 12395–12403 (2018).

36. Patterson, H. NHPatterson/napari-imsmicrolink: IMS MicroLink v0.1.7. (2022) doi:10.5281/ZENODO.6562052.

37. Luu, G. T. et al. TIMSCONVERT: a workflow to convert trapped ion mobility data to open data formats. Bioinformatics 38, 4046–4047 (2022).

38. Bemis, K. D. et al. Cardinal: an R package for statistical analysis of mass spectrometry-based imaging experiments. Bioinformatics 31, 2418–2420 (2015).

39. Sofroniew, N. et al. napari: a multi-dimensional image viewer for Python. GitHub Repository.

40. Sud, M. et al. LMSD: LIPID MAPS structure database. Nucleic Acids Res 35, D527–D532 (2007).

41. Li, Y., Wang, W., Jiang, T. & Yang, B. Aquaporins in urinary system. Adv Exp Med Biol 969, 131–148 (2017).

42. Rivenson, Y., de Haan, K., Dean Wallace, W. & Ozcan, A. Review Article Emerging Advances to Transform Histopathology Using Virtual Staining. (2020) doi:10.34133/2020/9647163.

43. Kaza, N., Ojaghi, A. & Robles, F. E. Virtual Staining, Segmentation, and Classification of Blood Smears for Label-Free Hematology Analysis. BME Front 2022, 1–14 (2022).

44. Ghahremani, P. et al. Deep learning-inferred multiplex immunofluorescence for immunohistochemical image quantification. Nature Machine Intelligence 2022 4:4 4, 401–412 (2022).

45. Li, Y. et al. Virtual histological staining of unlabeled autopsy tissue. Nature Communications 2024 15:1 15, 1–17 (2024).

46. Bai, B. et al. Deep learning-enabled virtual histological staining of biological samples. Light: Science & Applications 2023 12:1 12, 1–20 (2023).

47. Bejnordi, B. E. et al. Diagnostic Assessment of Deep Learning Algorithms for Detection of Lymph Node Metastases in Women With Breast Cancer. JAMA 318, 2199–2210 (2017).

48. Molenaar, M. R. et al. Increasing quantitation in spatial single-cell metabolomics by using fluorescence as ground truth. Front Mol Biosci 9, 1288 (2022).

49. Rappez, L. et al. SpaceM reveals metabolic states of single cells. Nature Methods 2021 18:7 18, 799–805 (2021).

50. Tideman, L. E. M. et al. Automated biomarker candidate discovery in imaging mass spectrometry data through spatially localized Shapley additive explanations. Anal Chim Acta 1177, 338522 (2021).

51. Guo, D., Bemis, K., Rawlins, C., Agar, J. & Vitek, O. Unsupervised segmentation of mass spectrometric ion images characterizes morphology of tissues. Bioinformatics 35, i208–i217 (2019).

52. Roy, A., Al-Bataineh, M. M. & Pastor-Soler, N. M. Collecting duct intercalated cell function and regulation. Clinical Journal of the American Society of Nephrology 10, 305–324 (2015).

53. Tokonami, N. et al. α-Ketoglutarate regulates acid-base balance through an intrarenal paracrine mechanism. J Clin Invest 123, 3166 (2013).

