## Supplemental Materials for "Automated Segmentation of Kidney Nephron Structures by Deep Learning Models on Label-free Autofluorescence Microscopy for Spatial Multi-omics Data Acquisition and Mining"

### Current affiliation: Aspect Analytics, C-Mine 12, 3600 Genk, Belgium.

---

#### Table of Contents

| | | |
|---|---|---|
| **Table S1** | Sample List……………………………………………………………………………………… | 2 |
| **Table S2** | Information on Fluorescence Filter Cubes used in Autofluorescence Experiments............ | 2 |
| **Table S3** | Information on Fluorescence Filter Cubes used in Multiplexed Immunofluorescence Experiments……………………………………………………………………………………….. | 3 |
| **Table S4** | Information on antibodies, filters, and exposure times for used in multiplex Immunofluorescence experiments. ……………………………………………………………….. | 3 |
| **Figure S1** | Functional tissue unit markers and multiplex immunofluorescence and autofluorescence microscopy images of human kidney tissues……………………………………………….. | 4 |
| **Figure S2** | Gross morphology (cortex, inner medulla, and outer medulla) differentiation with Aquaporin-1 and Uromodulin………………………………………………………………………. | 5 |
| **Figure S3** | Functional tissue unit segmentation modeling workflow…………………………………...… | 6 |
| **Table S5** | Training data instances count and statistics….…………………………………………….... | 6 |
| **Table S6** | Testing data instances count and statistics..…………………………………………………… | 6 |
| **Table S7** | Per sample global DICE metrics for each kidney functional tissue unit…………………….. | 7 |
| **Table S8** | Test set per sample instance-level metrics…………………………………………………….. | 7 |
| **Figure S4** | Workflow for spatially targeted transcriptomics data acquisition using gross morphological and functional tissue unit segmentation masks……………………………… | 9 |
| **Figure S5** | Aquaporin transcript expression in outer versus inner medulla……………………………… | 9 |

**Table S1.** Sample list

| # | Sample | Train | Test | Other | # | Sample | Train | Test | Other |
|---|---|---|---|---|---|---|---|---|---|
| **1** | VAN0005 | X | | | **20** | VAN0040 | X | X | |
| **2** | VAN0006 | X | X | | **21** | VAN0042 | X | X | |
| **3** | VAN0007 | X | | | **22** | VAN0043 | X | | |
| **4** | VAN0008 | X | | | **23** | VAN0044 | X | | |
| **5** | VAN0010 | X | | | **24** | VAN0045 | X | | |
| **6** | VAN0011 | X | | | **25** | VAN0046 | X | | |
| **7** | VAN0012 | X | | | **26** | VAN0047 | X | | |
| **8** | VAN0014 | X | | | **27** | VAN0048 | X | | |
| **9** | VAN0019 | X | | | **28** | VAN0049 | X | | |
| **10** | VAN0023 | X | X | | **29** | VAN0050 | X | | |
| **11** | VAN0024 | | | MALDI IMS | **30** | VAN0051 | X | X | |
| **12** | VAN0025 | X | | | **31** | VAN0052 | X | | |
| **13** | VAN0027 | X | | | **32** | VAN0053 | X | | |
| **14** | VAN0028 | X | | | **33** | VAN0054 | X | | GeoMX |
| **15** | VAN0029 | X | | | **34** | VAN0055 | X | | |
| **16** | VAN0030 | X | | | **35** | VAN0063 | X | X | |
| **17** | VAN0031 | X | | | **36** | KPMP-D1 | X | X | |
| **18** | VAN0033 | X | | | **37** | KPMP-D4 | X | | |
| **19** | VAN0038 | X | X | | | | | | |

**Table S2.** Information on fluorescence filter cubes used in autofluorescence experiments.

| Filter name | Filter set Zeiss ID | LED line wavelength (nm) | LED intensity setting | Excitation wavelength (nm) | Emission wavelength (nm) | Exposure time (ms) |
|---|---|---|---|---|---|---|
| DAPI | 90 HE DAPI / GFP / Cy3 / Cy5 | 385 | 90% | 375-395 | 410-440 | 20 |
| eGFP | 90 HE DAPI / GFP / Cy3 / Cy5 | 475 | 90% | 455-483 | 499-529 | 60 |
| Cy3 / DsRed | 90 HE DAPI / GFP / Cy3 / Cy5 | 567 | 90% | 545-565 | 579-604 | 250 |

AF images were collected using a Zeiss AxioScan.Z1 slide scanner with a 10x objective (Zeiss Plan-Apochromat 10x (NA=0.45) M27) with a Hamamatsu Orca Flash camera resulting in 16-bit images with 0.65 µm/px pixel spacing. Three common fluorescence filter wavelengths were used with a multi-band pass filter with illumination from a Zeiss Colibri7 LED light source.

**Table S3.** Information on fluorescence filter cubes used in multiplex immunofluorescence experiments.

| Filter name | Filter set Zeiss ID | LED line wavelength (nm) | LED intensity setting | Excitation wavelength (nm) | Emission wavelength (nm) |
|---|---|---|---|---|---|
| DAPI | 90 HE DAPI / GFP / Cy3 / Cy5 | 385 | 90% | 375-395 | 410-440 |
| eGFP | 90 HE DAPI / GFP / Cy3 / Cy5 | 475 | 90% | 455-483 | 499-529 |
| Cy5 | 90 HE DAPI / GFP / Cy3 / Cy5 | 630 | 90% | 620-643 | 659-759 |
| AF594 | 91 HE CFP / YFP / mCherry | 567 | 90% | 583-600 | 618-756 |

MxIF images were collected using a Zeiss AxioScan.Z1 (Carl Zeiss Microscopy, NY) slide scanner with a 10x objective (Zeiss Plan-Apochromat 10x (NA=0.45) M27) with a Hamamatsu Orca Flash camera resulting in 16-bit images with 0.65 µm/px pixel spacing. Four fluorescence filters were used. DAPI was always used to image Hoescht stained nuclei. Other filters were specific to fluorescently conjugated antibodies.

**Table S4.** Information on antibodies, filters, and exposure times for used in multiplex immunofluorescence experiments. Shaded table cells indicate markers used to train deep learning models.

| Cycle | Filter Name | Antibody | Structure | Exposure time (ms) |
|---|---|---|---|---|
| 1 | eGFP | NaCl Co-Transporter | Distal Tubules | 100 |
| 1 | Cy5 | Uromodulin | Thick Ascending Limb | 70 |
| 1 | AF594 | SLC5A12 | Proximal Tubules | 220 |
| 2 | eGFP | Collagen-IV (α5) | Basement membrane | 100 |
| 2 | Cy5 | CD31 | Endothelial cells | 150 |
| 2 | AF594 | Aquaporin-2 | Collecting Ducts | 180 |
| 3 | eGFP | Podocalyxin | Glomerular podocytes | 100 |
| 3 | Cy5 | Calbindin | Distal / Connecting Tubules | 150 |
| 3 | AF594 | α-Smooth Muscle Actin | Vascular smooth muscle cells | 180 |
| 4 | eGFP | Aquaporin-1 | Proximal Tubules / Descending thin limb | 100 |
| 4 | Cy5 | Collagen-IV (α1/2) | Basement membranes | 130 |

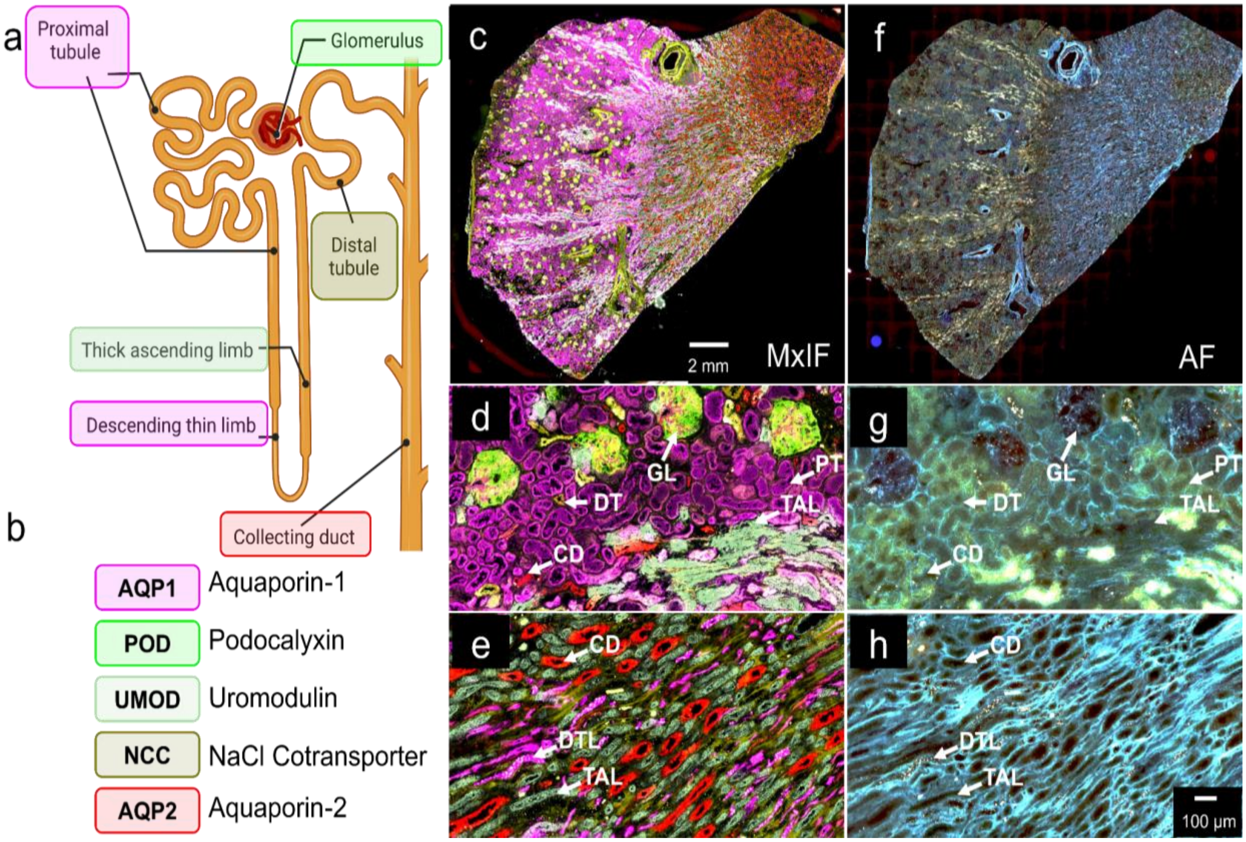


**Figure S1. Design of functional tissue unit (FTU) markers and comparison of multiplex immunofluorescence (MxIF) with label-free autofluorescence (AF) microscopy.** (a) Diagram of the nephron and its FTUs. (b) Color coding and abbreviations of FTU-specific antibodies used in MxIF. The same color coding persists in the MxIF microscopy images. *Note that there are only five distinct antibodies used to detect six FTUs, because AQP1 marks both the proximal convoluted tubule and the descending thin limb of the loop of Henle, the former in the kidney cortex and the latter in the medulla.* (c) Whole slide view of the combined MxIF data for all FTU markers. Higher magnification view of (d) the FTU MxIF image in the kidney cortex and (e) the FTU MxIF image in the kidney medulla. (f) Whole slide AF microscopy image of the same tissue section taken prior to MxIF. Higher magnification view of (g) the AF image in the kidney cortex and (h) the AF image in the kidney medulla. GL = Glomerulus; CD = Collecting Duct. DT = Distal Tubule. PT = Proximal Tubule. DTL = Descending Thin Limb. TAL= Thick Ascending Limb.

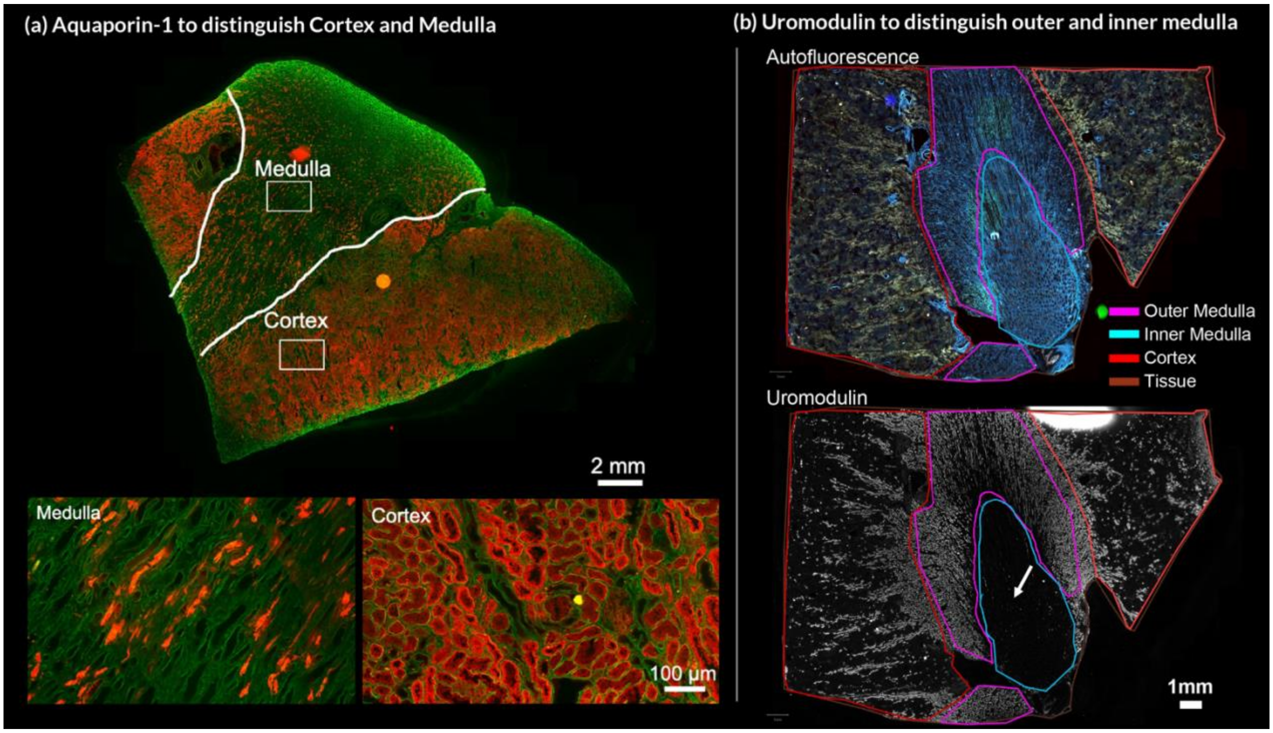


**Figure S2**. **Gross morphology (cortex, inner medulla, and outer medulla) differentiation with Aquaporin-1 and Uromodulin**. Comparison of Aquaporin-1 staining the human kidney medulla versus the cortex. (a) Top: Whole slide view of the sample with gross morphological areas and zoom-in areas highlighted. Bottom: Medulla and cortex comparison of morphology. *Green = Collagen-IV ($\alpha$1/2). Red = Aquaporin-1*. Autofluorescence and Uromodulin immunofluorescence to distinguish medullary zones. (b) Top: autofluorescence whole slide image with areas labeled by different colored polygons. Bottom: Uromodulin immunofluorescence with an arrow highlighting lack of Uromodulin signal in the inner medulla.

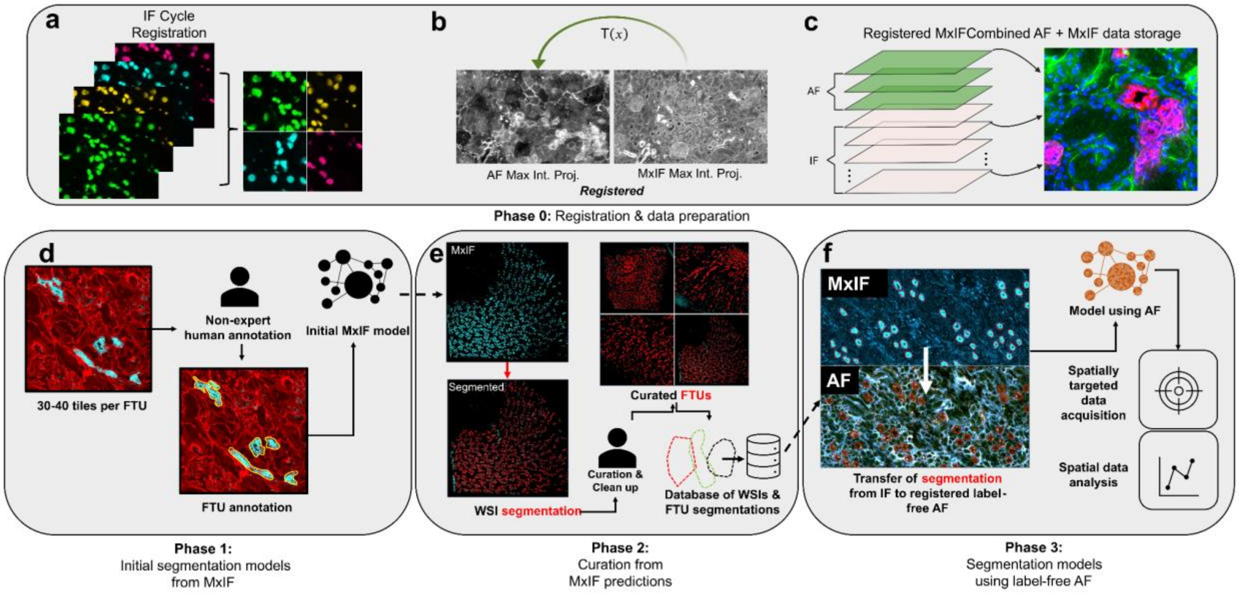


**Figure S3. Combined multiplex immunofluorescence (MxIF) and autofluorescence (AF) microscopy functional tissue unit (FTU) segmentation modeling workflow. (a-c)** *Phase 0*: Combination of MxIF images using the DAPI nuclear stain as a reference followed by registration of MxIF with AF from the same tissue section. Combined AF and MxIF images are stored in a single OME-TIFF so that annotations made in one modality can be used to subset the other. **(d)** *Phase 1*: Generation of initial MxIF based model to predict FTUs. The blue signal is AQP2 highlighting collecting ducts and the red signal is Collagen-IV (α1/2) highlighting FTU boundaries. **(e)** *Phase 2*: Application of IF model to whole slide MxIF images and curation of the segmented data into database of FTU segmentations. **(f)** *Phase 3*: Transfer of all segmentation data from the MxIF image to the AF image and training of deep learning models that use AF to segment FTUs. AQP2 = Aquaporin-2.

**Table S5.** Training data instances count and statistics.

| FTU | Number of WSIs in train data | Total train instances | Mean train instances | Median train instances | Std. Dev. train instances |
|---|---|---|---|---|---|
| Collecting Duct | 27 | 28661 | 1061.52 | 696 | 642.43 |
| Distal Tubule | 19 | 12558 | 660.95 | 603 | 329.27 |
| Descending Thin Limb | 17 | 14217 | 836.29 | 618 | 596.52 |
| Glomerulus | 27 | 5089 | 188.48 | 177 | 94.66 |
| Proximal Tubule | 24 | 39044 | 1626.83 | 1632 | 784.03 |
| Thick Ascending Limb | 19 | 12558 | 660.95 | 603 | 329.27 |

**Table S6.** Testing data instances count and statistics.

| FTU | Number of WSIs in test data | Total test instances | Mean test instances | Median test instances | Std. dev. test instances |
|---|---|---|---|---|---|
| Collecting Duct | 7 | 3188 | 455.43 | 218 | 417.85 |
| Distal Tubule | 7 | 1271 | 181.57 | 200 | 84.34 |
| Descending Thin Limb | 7 | 2725 | 454.17 | 406 | 250.83 |
| Glomerulus | 7 | 307 | 43.86 | 45 | 14.46 |
| Proximal Tubule | 7 | 10034 | 1433.43 | 1589 | 552.03 |
| Thick Ascending Limb | 7 | 7505 | 1072.14 | 995 | 601.11 |

**Table S7.** Per sample global DICE metrics for each kidney functional tissue unit.

| | Distal Tubule | Proximal Tubule | Collecting Duct | Glomerulus | Thick Ascending Limb | Descending Thin Limb |
|---|---|---|---|---|---|---|
| Sample ID | DICE | DICE | DICE | DICE | DICE | DICE |
| S1 | 0.8987 | 0.9535 | 0.9052 | 0.9877 | 0.8874 | 0.5090 |
| S2 | 0.8471 | 0.9294 | 0.9218 | 0.9716 | 0.8914 | 0.6658 |
| S3 | 0.8445 | 0.9488 | 0.8731 | 0.9645 | 0.8994 | 0.4951 |
| S4 | 0.8474 | 0.9580 | 0.8830 | 0.9698 | 0.9037 | 0.6582 |
| S5 | 0.8504 | 0.8912 | 0.8649 | 0.9486 | 0.8645 | 0.6646 |
| S6 | 0.8782 | 0.9579 | 0.8985 | 0.9643 | 0.9062 | 0.7065 |
| S7 | 0.8720 | 0.9532 | 0.9456 | 0.9221 | 0.9067 | 0.5242 |

**Table S8.** Test set per sample instance-level metrics.

| Sample ID | Distal Tubule | Proximal Tubule | Collecting Duct | Glomerulus | Thick Ascending Limb | Descending Thin Limb |
|---|---|---|---|---|---|---|
| | No. True Positive | No. True Positive | No. True Positive | No. True Positive | No. True Positive | No. True Positive |
| S1 | 188 | 1799 | 316 | 55 | 1306 | 102 |
| S2 | 152 | 1671 | 921 | 57 | 1904 | 440 |
| S3 | 89 | 1002 | 126 | 36 | 478 | 83 |
| S4 | 252 | 1749 | 979 | 44 | 853 | 209 |
| S5 | 192 | 1349 | 165 | 42 | 1219 | 130 |
| S6 | 93 | 818 | 136 | 33 | 445 | 110 |
| S7 | 170 | 1522 | 344 | 35 | 539 | 120 |

| Sample ID | Distal Tubule | Proximal Tubule | Collecting Duct | Glomerulus | Thick Ascending Limb | Descending Thin Limb |
|---|---|---|---|---|---|---|
| | No. False Positive | No. False Positive | No. False Positive | No. False Positive | No. False Positive | No. False Positive |
| S1 | 16 | 18 | 14 | 1 | 25 | 8 |
| S2 | 6 | 13 | 21 | 1 | 50 | 17 |
| S3 | 7 | 7 | 6 | 1 | 35 | 6 |
| S4 | 15 | 9 | 16 | 1 | 18 | 15 |
| S5 | 7 | 14 | 9 | 1 | 16 | 9 |
| S6 | 5 | 7 | 5 | 1 | 7 | 9 |
| S7 | 9 | 12 | 8 | 2 | 7 | 19 |

| Sample ID | Distal Tubule | Proximal Tubule | Collecting Duct | Glomerulus | Thick Ascending Limb | Descending Thin Limb |
|---|---|---|---|---|---|---|
| | Precision | Precision | Precision | Precision | Precision | Precision |
| S1 | 0.9216 | 0.9901 | 0.9576 | 0.9821 | 0.9812 | 0.9273 |
| S2 | 0.9620 | 0.9923 | 0.9777 | 0.9828 | 0.9744 | 0.9628 |
| S3 | 0.9271 | 0.9931 | 0.9545 | 0.9730 | 0.9318 | 0.9326 |
| S4 | 0.9438 | 0.9949 | 0.9839 | 0.9778 | 0.9793 | 0.9330 |
| S5 | 0.9648 | 0.9897 | 0.9483 | 0.9767 | 0.9870 | 0.9353 |
| S6 | 0.9490 | 0.9915 | 0.9645 | 0.9706 | 0.9845 | 0.9244 |
| S7 | 0.9497 | 0.9922 | 0.9773 | 0.9459 | 0.9872 | 0.8633 |

| Sample ID | Distal Tubule | Proximal Tubule | Collecting Duct | Glomerulus | Thick Ascending Limb | Descending Thin Limb |
|---|---|---|---|---|---|---|
| | Recall | Recall | Recall | Recall | Recall | Recall |
| S1 | 0.9447 | 0.9740 | 0.8977 | 0.9649 | 0.9409 | 0.2400 |
| S2 | 0.7835 | 0.8631 | 0.9228 | 0.9500 | 0.9292 | 0.4900 |
| S3 | 0.8165 | 0.9570 | 0.9197 | 0.9730 | 0.9775 | 0.2562 |
| S4 | 0.7975 | 0.9002 | 0.8998 | 0.9167 | 0.9094 | 0.4172 |
| S5 | 0.9366 | 0.8765 | 0.8730 | 0.9333 | 0.9165 | 0.5532 |
| S6 | 0.8774 | 0.9446 | 0.8553 | 0.9167 | 0.9408 | 0.5314 |
| S7 | 0.8523 | 0.9112 | 0.8823 | 0.9225 | 0.9113 | 0.4889 |

| Sample ID | Distal Tubule | Proximal Tubule | Collecting Duct | Glomerulus | Thick Ascending Limb | Descending Thin Limb |
|---|---|---|---|---|---|---|
| | F1-Score | F1-Score | F1-Score | F1-Score | F1-Score | F1-Score |
| S1 | 0.9330 | 0.9820 | 0.9267 | 0.9735 | 0.9606 | 0.3813 |
| S2 | 0.8636 | 0.9232 | 0.9495 | 0.9661 | 0.9513 | 0.6494 |
| S3 | 0.8683 | 0.9747 | 0.9368 | 0.9730 | 0.9541 | 0.4019 |
| S4 | 0.8645 | 0.9451 | 0.9400 | 0.9462 | 0.9431 | 0.5766 |
| S5 | 0.9505 | 0.9297 | 0.9091 | 0.9545 | 0.9505 | 0.6952 |
| S6 | 0.9118 | 0.9675 | 0.9067 | 0.9429 | 0.9622 | 0.6748 |
| S7 | 0.8984 | 0.9500 | 0.9274 | 0.9341 | 0.9477 | 0.6243 |

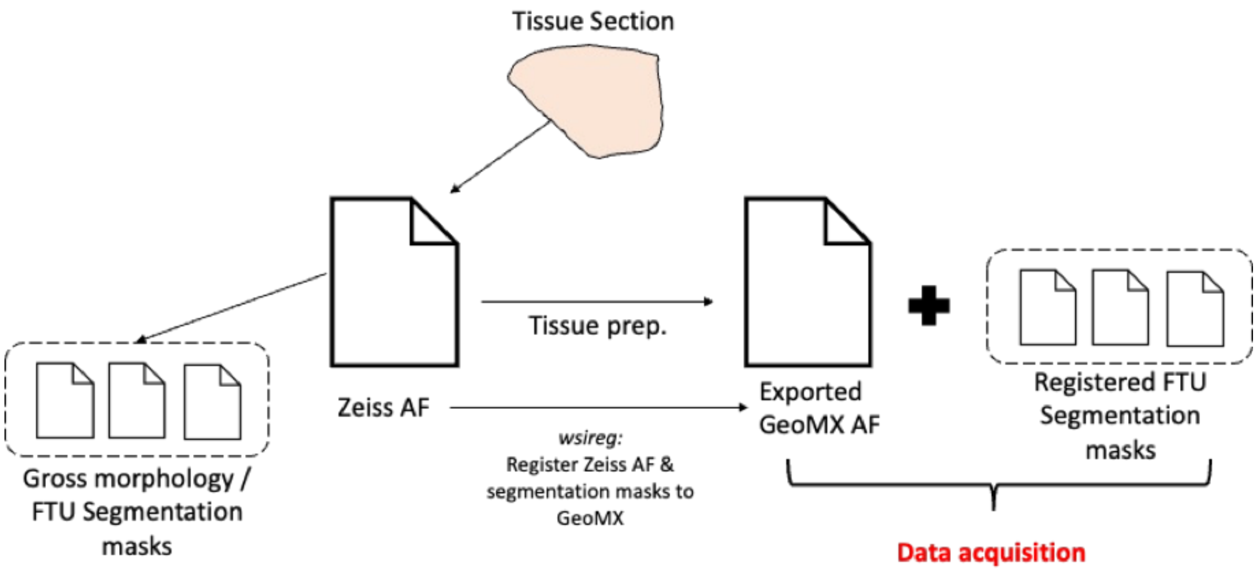


**Figure S4.** Workflow for spatially targeted transcriptomics data acquisition using gross morphological and functional tissue unit segmentation masks. AF = autofluorescence microscopy. FTU = Functional tissue unit. GeoMX = Nanostring GeoMX Digital Spatial Profiler instrument

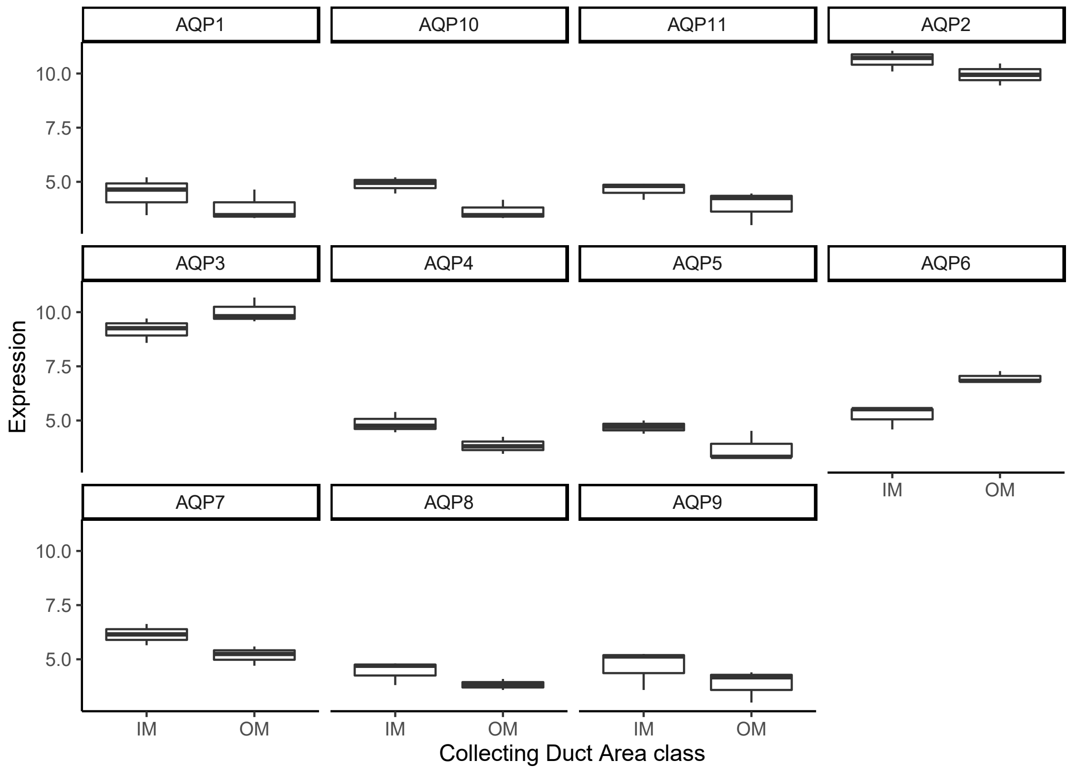


**Figure S5.** Aquaporin transcript expression in outer versus inner medulla. IM= Inner medulla. OM = Outer medulla.
